# Maternal immune activation during the lactational period alters offspring behavior, reproductive development, and immune function in mice

**DOI:** 10.1101/2025.05.26.656156

**Authors:** Jailyn A. Merengueli, Amanda C. Kentner

**Author notes:** Corresponding author: Amanda Kentner Office.

## Abstract

Exposure to infection during early life can lead to lasting neurodevelopmental changes. Animal models of maternal immune activation (MIA) typically assess neurobehavioral alterations in offspring following a prenatal inflammatory challenge. However, MIA effects on offspring can extend to challenges that occur during the lactational period. In the present study, we adapted previous methods focused on rats and challenged nursing C57BL/6J mouse mothers on postnatal day (P)8 with either the bacterial mimetic lipopolysaccharide (LPS; 250 μg/kg, i.p.) or saline (control, i.p.). By exposing only the mother to LPS, this modeled a postpartum infection in the dam. Similar to the rat model, lactational MIA did not detrimentally alter maternal care but induced displays of maternal sickness, as expected. While neonatal offspring behaviors (e.g., huddling, ultrasonic vocalizations, negative geotaxis) were unaffected, significant effects of lactational MIA emerged in juvenile (e.g., social preference, accelerated reproductive milestones) and adult (e.g., mechanical allodynia, prepulse inhibition) offspring. In a separate set of animals, the developmental programming potential of lactational MIA on immune function was evident following a “second hit” LPS challenge in adulthood (e.g., altered plasma concentrations of interleukin-6 and leukocytes, including neutrophils, and lymphocytes). These findings confirm the generalizability of the lactational MIA model across species and highlight the importance of supporting caregiver health and wellness across the critical nursing period.

## Introduction

The perinatal period represents a critical window of neurodevelopmental plasticity, during which environmental experiences can exert lasting effects on brain structure and function. A wide body of literature has established that maternal immune activation (MIA) during gestation can alter offspring neurodevelopment and confer increased risk for a range of neuropsychiatric disorders, including autism spectrum disorder and schizophrenia (Estes & McAllister, 2016; Knuesel et al., 2014; Gumusoglu et al., 2019). In the animal laboratory, systemic administration of immune mimetics such as lipopolysaccharide (LPS) or polyinosinic:polycytidylic acid (poly I:C) to pregnant rodent dams’ results in maternal cytokine (e.g., IL-6, IL-1β, TNF-α) and hormone (e.g., corticosterone) release, and placental inflammation and stress responses. This is in addition to changes in other processes known to impact key pathways including microglial maturation, synaptogenesis, and hypothalamic-pituitary-adrenal (HPA) axis programming, which also account for altered fetal brain development, (Martz et al., 2024; Knuesel et al., 2014; Bilbo & Schwarz, 2012; Kentner et al., 2019; Gumusoglu et al., 2019; Cattane et al., 2022; Otero & Antonson, 2022).

Much less is known about the consequences of immune challenges that occur after birth, when the maternal-offspring dyad is still connected through nursing. During this time, offspring are influenced via “*lactocrine*” signaling (e.g., inflammatory cytokines and glucocorticoids in milk; Sullivan et al., 2011; Bartol et al., 2013; Hinde et al., 2015; de Weerth, et al., 2023; Ziomkiewicz et al., 2021), behavioral cues (e.g., maternal care; Champagne et al., 2003; Francis, et al., 2002; van Hasselt et al., 2012), and chemosensory exposure (e.g., maternal scent/pheromonal signals; Shaal et al., 2003); each of these critical communication pathways can be modified via immune challenges (DeRosa et al., 2022; Vilela et al., 2013; Ling et al., 2010; Wijenayake et al., 2023; Macrae et al., 2015a; Arakawa et al., 2011). Moreover, the nursing period is increasingly recognized as a time of continued neuroimmune vulnerability, during which direct disruptions to either pups (Shanks et al., 1995; Walker et al., 2009; Bilbo et al., 2005; Spencer et al., 2006) or dams (Vilela et al., 2013; Nascimento et al., 2015; DeRosa et al., 2022) may exert immediate, and/or latent effects on offspring development.

To explore these postnatal dynamics, traditional rodent MIA models have been adapted to the lactational period (Vilela et al., 2013; Nascimento et al., 2015; DeRosa et al., 2022), demonstrating that maternal LPS exposure in the early postnatal days can lead to altered juvenile and adult offspring behavior (Nascimento et al., 2015; DeRosa et al., 2022). Notably, these behavioral alterations can occur without adversely disrupting maternal care (Aubert et al., 1997; DeRosa et al., 2022; Nascimento et al., 2015) and despite the fact that the offspring are not directly exposed to LPS themselves (e.g., LPS did not cross through the milk after the postpartum immune challenge), underscoring the potential for other indirect maternal mechanisms (DeRosa et al., 2022). In our prior work using rats, we reported that lactational MIA induced long-term changes in offspring behaviors relevant to social thermoregulation, tactile sensitivity, and sensorimotor gating. However, the generalizability of this lactational MIA model across species remains unknown. This is a critical limitation, given the field’s increasing emphasis on rigor, reproducibility, and the translational impact of preclinical research (Collins & Tabak, 2014; Kentner et al., 2018). Indeed, in humans, postpartum infections are common, yet the effects of maternal illness on nursing practices and infant outcomes are understudied (PRGLAC, 2018). Breastfeeding is still recommended during most maternal illnesses, as transmission of common pathogens through breastmilk is rare (CDC, 2025; Yeo et al., 2022). However, both illness and other types of stressors can impact milk quality, and more research is needed to fully understand the clinical implications of these changes and how supportive interventions may be implemented (Wijenayake et al., 2023).

Cross-species replication is essential for identifying conserved pathways in addition to improving the robustness of model systems and behavioral constructs (Gulinello et al., 2020; van Goethem et al., 2012; Geyer et al., 2008; Kafkafi et al., 2018). C57BL/6 mouse substrains in particular represent a foundational species in neuroscience due to the extensive characterization and widespread use of these mice in behavioral and genetic studies (Bryant, 2011). Challenging the robustness of the lactational MIA model in C57BL/6 mice is therefore a reasonable choice to 1) test whether the previously observed behavioral phenotypes in rats can be replicated, and 2) determine whether general neuroimmune mechanisms are conserved across species.

Consequently, we aimed to evaluate the cross-species generalizability of the lactational MIA model by administering LPS to nursing C57BL/6J mouse dams on postnatal day 8 (Clancy et al., 2007), a period of active synaptogenesis and myelination in developing offspring (Semple et al., 2013). A range of offspring behaviors were assessed across different developmental stages, as was reproductive timing. In adulthood, we further investigated the long-term impact of lactational MIA on immune responsiveness to a subsequent inflammatory insult. Indeed, reprogramming by the earlier MIA, or “first hit” challenge may result in either increased vulnerability or adaptive resilience, depending on context and timing of a “second hit” challenge (Giovanoli et al., 2013; Calcia et al., 2016: Maynard et al., 2001). To explore this, adult offspring were administered either LPS or saline after which we measured plasma corticosterone (Walker et al., 2009), interleukin (IL)-6 (Ellis et al., 2005; Kentner at al., 2010), and leukocyte biomarkers (Sailhamer et al., 2010) to assess stress and immune reactivity. This design allowed us to test whether lactational MIA acts as a developmental programming event, impacting susceptibility to later inflammatory or stress-related disorders (Bilbo et al., 2005; Bilbo et al., 2008; Ellis et al., 2005; Ellis et al., 2006; Walker et al., 2009; Spencer et al., 2011).

## Materials and Methods

### Animals and Husbandry

C57BL/6J mice (Jackson Laboratory, Bar Harbor, ME) were housed in polycarbonate cages (7 1⁄2” x 11 1⁄2” x 5”) and bred in groups of one male to two females (at 21°C on a 12 h light/dark cycle, 0700–1900 light) with ad libitum access to food and water. Animals had access to one chew bone and Bed-r’Nests® (ScottPharma Solutions, Marlborough MA). Pregnancy was confirmed by observations of a sustained diestrus phase (observed from evaluation of vaginal samples; Marcondes et al., 2002; Zhao et al., 2021), in addition to continued weight gain during the later phase of gestation. Females were separated and singly housed on ∼gestational day 15 with extra Bed-r’Nest® materials and the chew bone was removed and replaced at weaning. Day of birth was identified as postnatal day (P)0 and litters were standardized to n = 8 pups per litter on P5, ensuring an equal balance between male and female pups whenever possible.

To adapt our lactational MIA model (DeRosa et al., 2022) from rats to mice, dams were administered either 250µg/kg lipopolysaccharide (LPS, Escherichia coli, serotype 026:B6; L-3755, i.p; Weil et al., 2006; Aubert et al., 1997) or equalvolume saline (i.p) on P8 (Clancy et al., 2007; n = 9 litters/group). On P21, offspring were ear clipped to designate their treatment group and weaned into same sex pairs, housed as one LPS and one saline animal per cage. Investigators were blinded to the group designations. Only one male and one female per litter were used on each group measure throughout the study. The general experimental timeline is displayed in **Figure 1**.

**Figure 1.**
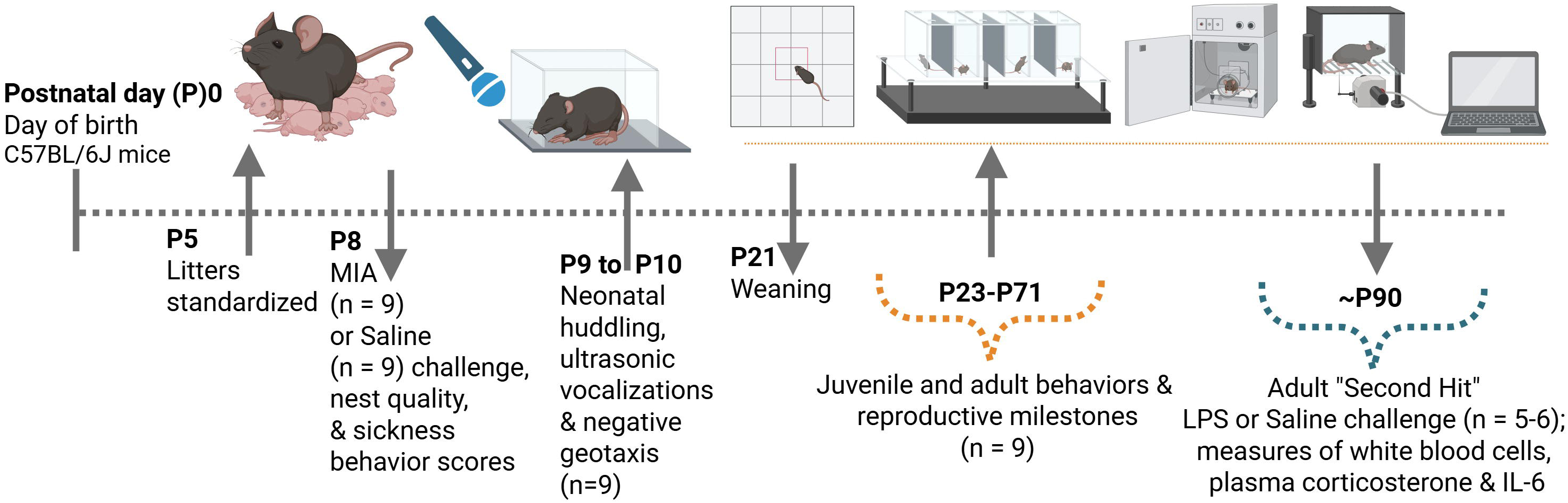
Experimental Timeline. Created in BioRender. Kentner, A. (2025) https://BioRender.com/86qqgzg

The MCPHS University Institutional Care and Use Committee approved all procedures described, which were carried out in compliance with the recommendations outlined by the Guide for the Care and Use of Laboratory Animals of the National Institutes of Health.

### Physiological and Behavioral Measures

#### Maternal sickness behaviors and nest construction quality

Building a nest from fresh materials requires a high level of activity and engagement, which is an indicator of maternal motivation. Our mice tend to construct dome-shaped nests, or simply cover the litter with the nesting materials, obscuring maternal-pup interactions. Therefore, as a proxy of maternal care quality, and a way to dissociate the effects of immune activation from variations in care, we evaluated nest structures using a 5-point Likert scale, adapted from Gerfen et al., (2006; Zhao et al., 2021) in which 0 = no nest; 1 = flat nest; 2 = a slightly cupped shape but less than 1/2 of a dome shape; 3 = taller than 1/2 of a dome shape, but not fully enclosed and 4 = fully enclosed nest/dome. Maternal sickness behaviors were scored based on the absence (score = 0) or presence (score = 1) of illness indicators which were summed into a composite score (range from 0 to 4). Behaviors included piloerection (fur standing on end), ptosis (drooping upper eyelid), lethargy (lack of movement/low energy), and huddling behavior (curled body posture). On the morning of P8, baseline evaluations assessed the presence of maternal sickness behaviors and nest structure quality, prior to MIA induction. At the point of the P8 MIA challenge, nests were removed from the cage and fresh nesting materials were provided. In addition to the baseline evaluation, nest quality and the magnitude of sickness behaviors were again evaluated at 2-, 3-, 6-, and 24-hours post MIA challenge.

#### Neonatal huddling (P9 & P10)

To evaluate social thermoregulation behaviors, each full litter completed a neonatal huddling test on P9 and P10 as previously described (Naskar et al., 2019; DeRosa et al., 2022). Lactational MIA has previously reduced huddling behavior in neonatal rats (DeRosa et al., 2022). Pups were placed at equal distances from each other along the perimeter of a test arena (40 cm x 40 cm). Pups were videorecorded for 10 minutes. Using one video frame for every 30 s of the video, the total average number of pup clusters (a cluster is defined as two or more pups in physical contact) across the test period was calculated.

#### Negative geotaxis (P10)

This sensorimotor test evaluates the natural tendency to turn oneself facing up towards a slope, in response to vestibular cues of gravity, when placed facing down a slope (Ruhela et l., 2019; Feather-Schussler et al., 2016). Pups were placed upside down on a 45° inclined plane and time (s) to rotate 180° was recorded. Trials terminated after 60 s.

Each animal underwent three consecutive trials and the mean time (s) to rotate was calculated. This test was added to evaluate other aspects of sensorimotor functioning that may be disrupted by lactational MIA.

#### Ultrasonic vocalizations (USVs; P10)

Neonatal USVs, modulated by social cues, are disrupted in the gestational MIA model (Martz et al., 2024). Here, we tested whether that disruption extends to the lactational challenge model. Pups were individually removed from their litters and housed in a small cage lined with Nestlets®. To measure isolation-induced USVs, the cage was placed into a sound attenuation chamber and isolation calls were recorded from an ultrasonic microphone (UltraSoundGate 116–200), located 20 cm from the floor, for 5-minutes. The USV microphone was connected to a PC and the software program Audacity was used to record vocalizations at 192 kHz, capturing a range of USVs. Vocalizations were analyzed by automated Live Mouse Tracking (LMT) USV analysis software (de Chaumont et al., 2019), to determine the total number of calls emitted from each vocalization recording.

#### Reproductive milestones

Following gestational MIA (Zhao et al., 2022) and in other models of early-life stress (Grassi-Oliveira et al., 2016; Granata et al., 2024), the initiation of some developmental milestones such as puberty are accelerated. Therefore, starting on P23, male and female mice underwent daily evaluations to confirm age of complete balanopreputial separation or vaginal opening (closed or complete; Grassi-Oliveira et al., 2016; Granata et al., 2024; Zhao et al., 2022).

#### Open Field Test (P23 and P70)

Prior to the social preference test (see below) mice were habituated to the open field test arena (40 cm x 40 cm x 28 cm) and video recorded for 5 min (Martz et al., 2022). Automated behavioral monitoring software (Any-maze, Wood Dale, IL) evaluated open field behavior by determining time (seconds) spent in each of the perimeter and center of the test arena, which was then converted as a percentage of time spent in the center of the open field [total time in center/total time in perimeter + total time in center) × 100] as an index of anxiety-like behavior.

#### Social Preference Index (P23 and P70)

Offspring social interest is significantly reduced following gestational MIA challenge (Martz et al., 2022). To determine if we could observe this effect in our lactational mouse model, immediately following evaluation of locomotor activity, a five-minute social preference test was videorecorded. The video was scored using automated behavioral monitoring software (Any-maze, Wood Dale, IL). The test consisted of two cleaned wire cups placed on either end of the arena. One cup held a novel untreated mouse of the same sex, age, size, and strain and the other cup held a novel object. The location of novel mice and novel objects was interchanged between trials. Experimental mice were allowed to freely explore the arena during the test period. A social preference index was calculated by the equation ([time spent with the mouse] / [time spent with the inanimate object + time spent with the mouse]) − 0.5; Scarborough et al., 2020).

#### The von Frey Test (P33, P45, P71)

Following a 30-minute habituation period, mechanical allodynia was evaluated using a pressure-meter comprised of a hand-held force transducer fitted with a 0.7 mm^2^ polypropylene tip (electronic von Frey anesthesiometer, IITC, Inc, Life Science Instruments, Woodland Hills, CA, USA). The polypropylene tip was applied to the animal’s left hind paw with an increasing force until a flexion reflex occurred. The threshold (grams) of the applied weight was automatically recorded by the electronic pressure-meter when the animal withdrew its paw. The average of three test trials was calculated as the mechanical allodynia threshold (Yan and Kentner, 2017).

#### Prepulse inhibition (PPI; P70)

San Diego Instruments chambers (San Diego, CA) were used to evaluate prepulse inhibition (PPI) of the acoustic startle reflex. Following a published protocol (Zhao et al., 2022), the trial consisted of pulse-alone, prepulse-plus-pulse and prepulse-alone trials, as well as no-stimulus trials in which only a background noise of 65 dB was presented. During stimulus trials, animals were exposed to a 40 ms pulse of 120 dB white noise with or without the presentation of a prepulse. Prepulse intensity consisted of one of five intensity types: 69, 73, 77, 81, 85 dB corresponding to 4, 8, 12, 16, and 20 dB above the background noise that was administered prior to the 120 dB white noise. A 100 ms interval was used between the prepulse and the pulse during the prepulse-plus-pulse trials. A 20-ms burst of 5 different prepulses was administered during the prepulse-alone trials. Each session was initiated by 6 pulse-alone trials. Then, each prepulse-plus-pulse, prepulse-alone, pulse-alone and no-stimulus trial was presented 12 times in a pseudorandom order. The session ended with 6 successive pulse-alone trials. The average interval between successive trials (ITI) was 15 ± 5 sec. For each animal and at each of the five possible prepulse intensities, the % PPI for each prepulse intensity was calculated using the following formula: 1-(mean reactivity on prepulse-plus-pulse trials / mean reactivity on pulse alone trials) × 1/100 (Giovanoli et al. 2014).

### Adult Treatment and White Blood Cell Analysis

Previous studies demonstrate that early life stress manipulations can modify immune reactivity in later life (Neveu et al., 1994; Clark et al., 2018; Makinson et al., 2017;Nascimento et al., 2015). Given the importance of white blood cells to overall health and immune functioning (Sailhamer et al., 2010; Nicholson, 2016), we aimed to determine whether lactational MIA challenge modified the concentrations of these cells, either in the absence or presence of a second LPS challenge. On ∼P90, siblings of mice used in the behavioral studies were subdivided to receive an i.p injection of either LPS (50 μg/kg) or an equivalent volume of pyrogen-free saline. An additional two LPS and two saline litters were added to achieve the required animal numbers for the second hit challenge component of the study.

Three hours following the adult LPS challenge, animals were deeply anesthetized with isoflurane, decapitated, and trunk blood collected into EDTA blood collection tubes (BD, VT365974). Whole blood was analyzed using a HEMAVET automated multi-species hematology analyzer (M-950HV, Drew Scientific) to obtain white blood cell measurements (n = 5-6; two samples were lost due to blood clotting). Measures included the number of leukocytes in the evaluated volume (20µl) of whole blood directly measured, in addition to the absolute number of leukocytes that were neutrophils, lymphocytes, monocytes, eosinophils, or basophils, directly measured (10^9^ cells/liter) using the International System of Units. The remaining sample was centrifuged, and plasma aliquots were stored at 75°C until processing.

### Plasma Corticosterone and Interleukin (IL)-6 ELISAs

Second hit exposures to stress can influence plasma corticosterone and cytokine levels (Walker et al., 2009; Ellis, et al., 2005), affecting immune function. Therefore, we also evaluated this hormone and the cytokine IL-6 in blood plasma of adult animals (n = 5) three hours post LPS or saline challenge (as described above). Plasma corticosterone was measured in duplicate by ELISA according to the small sample assay protocol in the manufacturer’s instructions (ADI-900-097, Enzo Life Sciences, Farmingdale, NY, USA). The minimum detectable concentration was 26.99 pg/ml, and the intra- and inter-assay coefficients of variation were 6.6% and 7.8%, respectively. IL-6 was similarly measured following manufacturer’s instructions (Invitrogen, BMS603-2). Here, the minimum detectable concentration was 6.5 pg/ml, and the intra- and inter-assay coefficients of variation were 5.0% and 8.9%, respectively.

### Statistics

Statistical Software for the Social Sciences (SPSS) was used to perform statistical analyses. While we include both males and females in our study, separate analyses were run for each sex. Non-parametric Friedman’s analyses (version of the repeated measures ANOVA) were used to evaluate maternal nest construction quality and the magnitude of sickness behaviors. These were followed by Kruskal-Wallis post hocs, using an adjusted p-value of 0.01 to indicate significance, accounting for multiple comparisons. One-way (LPS vs Saline) repeated measures ANOVAs were used to evaluate huddling behavior, %PPI, and mechanical allodynia. Age of balano-preputial separation and vaginal opening were each assessed using the Log-rank (Mantel Cox) test and plotted as a survival curve. All other behavioral tests were evaluated by t-tests unless the data were skewed (Shapiro-Wilks), in which case non-parametric Kruskal-Wallis tests were employed (**Green et al., 2005**). Two-way ANOVAs (lactational MIA treatment X adult LPS treatment) were used in the analyses of the adult second-hit white blood cell and ELISA data.

These were followed by t-test post hocs, in the occurrence of fewer than three groups. As expected, some samples were below the level of assay detection (e.g., no inflammation was detected in Saline-Saline treated animals for the IL-6 assay); therefore, standard pharmacological approaches were applied, such as replacing undetectable values with zero, to enable statistical evaluation (Barnett et al., 2021). The partial eta-squared (*np*^2^) is reported as an index of effect size (Miles & Shevlin, 2001). Data are expressed as mean ± SEM.

## Results

### Lactational MIA induced sickness behaviors in mouse dams with only a modest and transient impact on maternal nest construction quality

As anticipated, lactational MIA challenge significantly increased displays of maternal illness across the evaluation period (MIA: *X*^2^(4) = 12.119, p = 0.016; Saline: p>0.05; **Figure 2A**). Maternal sickness scores were significantly higher 3- and 6-hours post MIA challenge compared to saline controls (3-hours: *X*^2^(1) = 6.548, p = 0.011; 6-hours: *X*^2^(1) = 7.440, p = 0.006). In contrast, nest construction quality varied across time in both LPS (*X*^2^(1) = 29.377, p = 0.001) and Saline (*X*^2^(1) = 14.000, p = 0.007) dams suggested that there were either no or minimal group differences (**Figure 2B**). Indeed, Kruskal-Wallis analysis revealed only an incremental and transient difference in nest quality score 2-hours post MIA (*X*^2^(1) = 5.497, p = 0.019).

**Figure 2.**
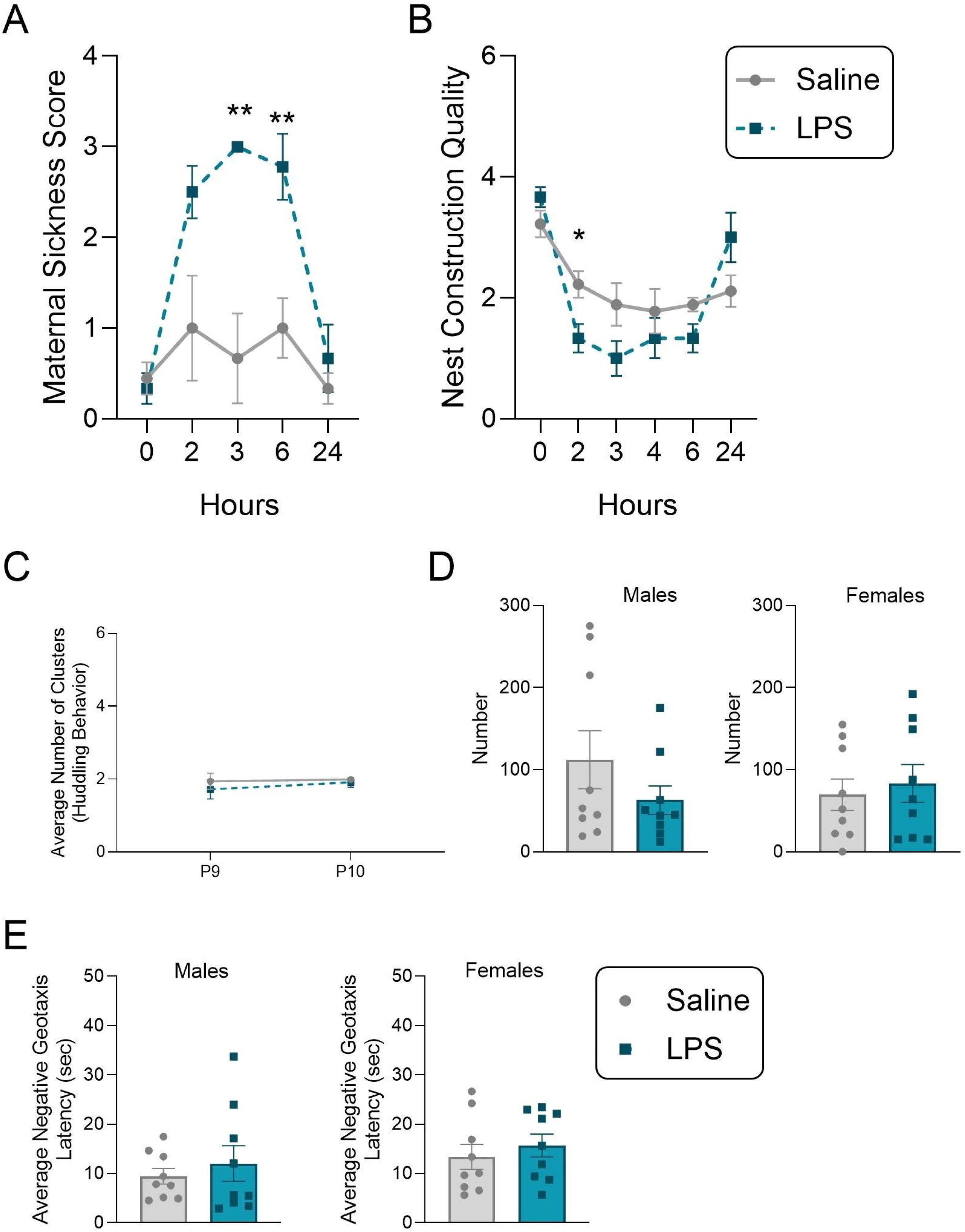
Maternal and neonatal outcomes following lactational maternal immune activation (MIA) in mice. A) Mouse dams treated with LPS on postnatal day (P) 8 had higher sickness scores than dams treated with saline, validating MIA. B) LPS dams had a modest and transient decrease in nest quality on the day of MIA challenge, suggesting that maternal care was not strongly compromised. C) Neonatal huddling behavior was not affected by MIA across P9 and P10. Neonatal behaviors including D) the total number of calls emitted during the isolation induced ultrasonic vocalization test and E) negative geotaxis were not affected by MIA (males left-side, females right-side). Saline: grey circles; lipopolysaccharide (LPS): blue squares. Data expressed as mean ± SEM, n = 9 litters/group. *p <0.05; **p< 0.01, Saline versus LPS.

### Lactational MIA did not affect neonatal offspring behavior

There were no significant effects of lactational MIA on neonatal huddling behavior, negative geotaxis latency, or the total number of USVs emitted by male or female offspring (p>0.05; **Figure 2C-E**).

### Lactational MIA was associated with altered juvenile and adult offspring behaviors and accelerated reproductive development

With respect to reproductive milestones, female MIA mice displayed accelerated puberty (t(16) = −2.530, p = 0.011, *np^2^* = 0.022; **Figure 3A**). Specifically, the Log-rank (Mantel Cox) test survival curve indicated that female MIA offspring displayed full vaginal openings approximately one day earlier than saline controls (*X^2^*(1) = 4.955, p = 0.026; **Figure 3B**). Juvenile male MIA offspring demonstrated an increased social preference index (independent t-test: t(16) = 2.2883, p = 0.011, *np^2^*= 0.0342) and female MIA offspring had a reduced social preference (independent t-test: t(16) = −2.20, p = 0.041, *np^2^* = 0.235) compared to their saline controls (**Figure 3C**). Percent time in the center of the open field was not affected in either juvenile male or female offspring (p>0.05, **Figure 3D**), suggesting that differences in social preference in response to MIA were not mediated by anxiety-like behavior. In adulthood, neither social preference nor percent time in the center of the open field differed between lactational MIA and saline offspring (p>0.05; **Supplemental Figure 1A,B**).

**Figure 3.**
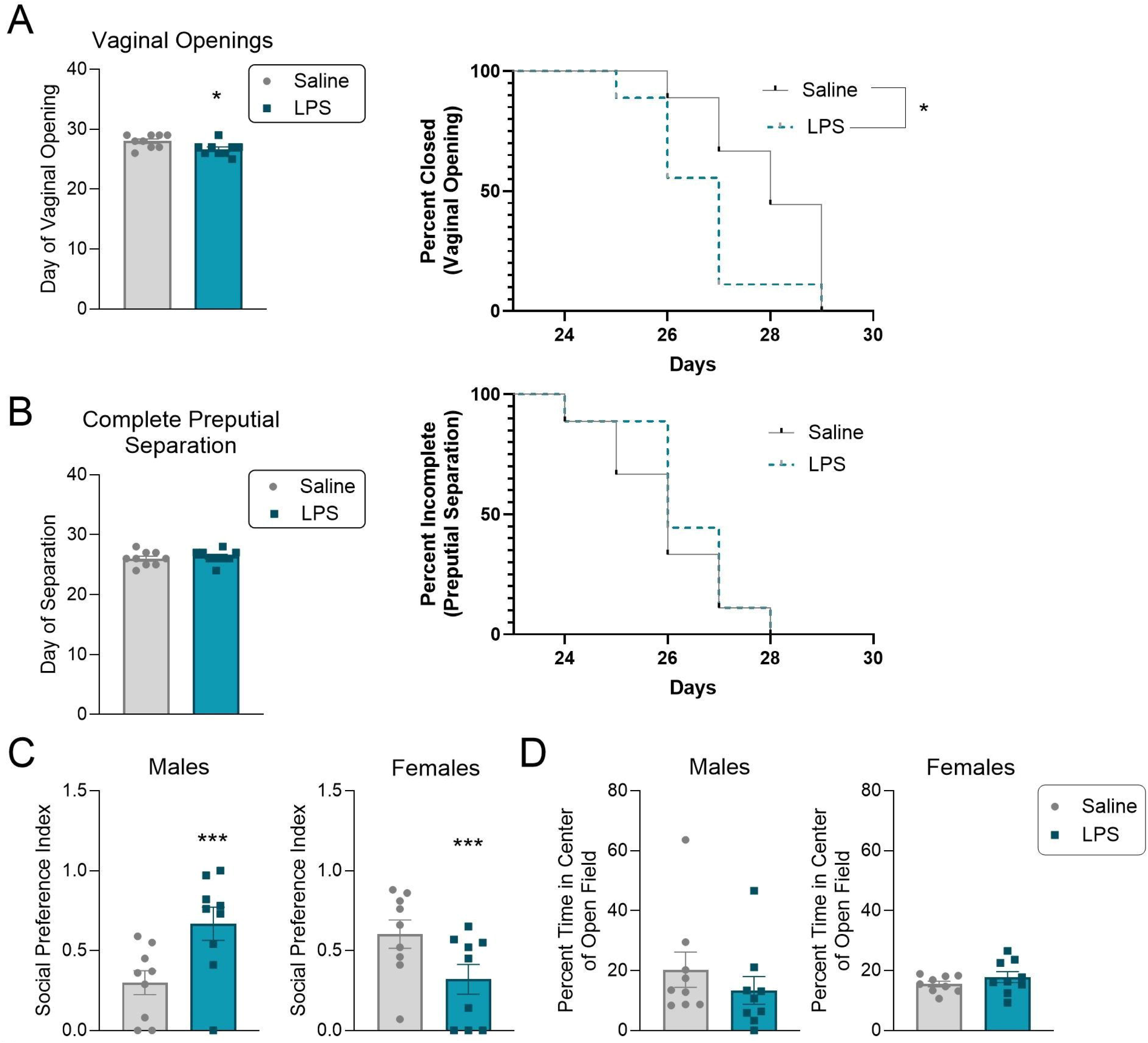
Juvenile offspring behaviors and reproductive milestones were affected by lactational maternal immune activation (MIA). A) The day of vaginal opening was accelerated in female LPS compared to saline female mice (top) while B) male preputial separation was not affected (independent t-tests left graphs; Log-rank (Mantel Cox) tests right graphs). Lactational MIA C) increased social preference in males (left-side) while decreasing it in females (right-side). Since D) percent time in the center of the open field was not affected in either male or female offspring, this suggest the social changes were not mediated by anxiety-like behavior. E) Saline: grey circles; lipopolysaccharide (LPS): blue squares. Data expressed as mean ± SEM, n = 9 litters/group. *p<0.05; **p<0.01; ***p<0.001, Saline versus LPS.

Repeated measures ANOVA confirmed that percent prepulse inhibition (%PPI) of the acoustic startle response increased as decibel (dB) level increased, as expected in adult animals (males: F(4, 64) = 46.552, p = 0.001, *np^2^* = 0.744; females: F(4, 64) = 21.133, p = 0.001, *np^2^* = 0.569; **Figure 4A**). Lactational MIA did not affect %PPI sensorimotor gating in adult female offspring (p>0.05, right-side graph). Meanwhile, adult male MIA mice (left-side graph) had reduced %PPI (Treatment by Time (dB): F(4, 64) = 4.070, p = 4.070, p = 0.005, *np^2^* = 0.203; post hoc independent t-test at 77dB = t(16) = −3.357, p = 0.004), as similarly seen in lactational MIA rats (DeRosa et al., 2022). A one-tailed independent t-test showed a significant decrease in mean %PPI for male MIA offspring compared to saline (t(16) = −2.029, p = 0.030, *np^2^* = 0.205; females: p>0.05; **Figure 4B**). Lactational MIA decreased mechanical allodynia thresholds (grams) in MIA male offspring while increasing it in females, as measured in the von Frey test (males: F(1, 13) = 4.772, p = 0.048, *np^2^* = 0.269; females: F(1, 13) = 5.008, p = 0.043, *np^2^* = 0.279; **Figure 4C**).

**Figure 4.**
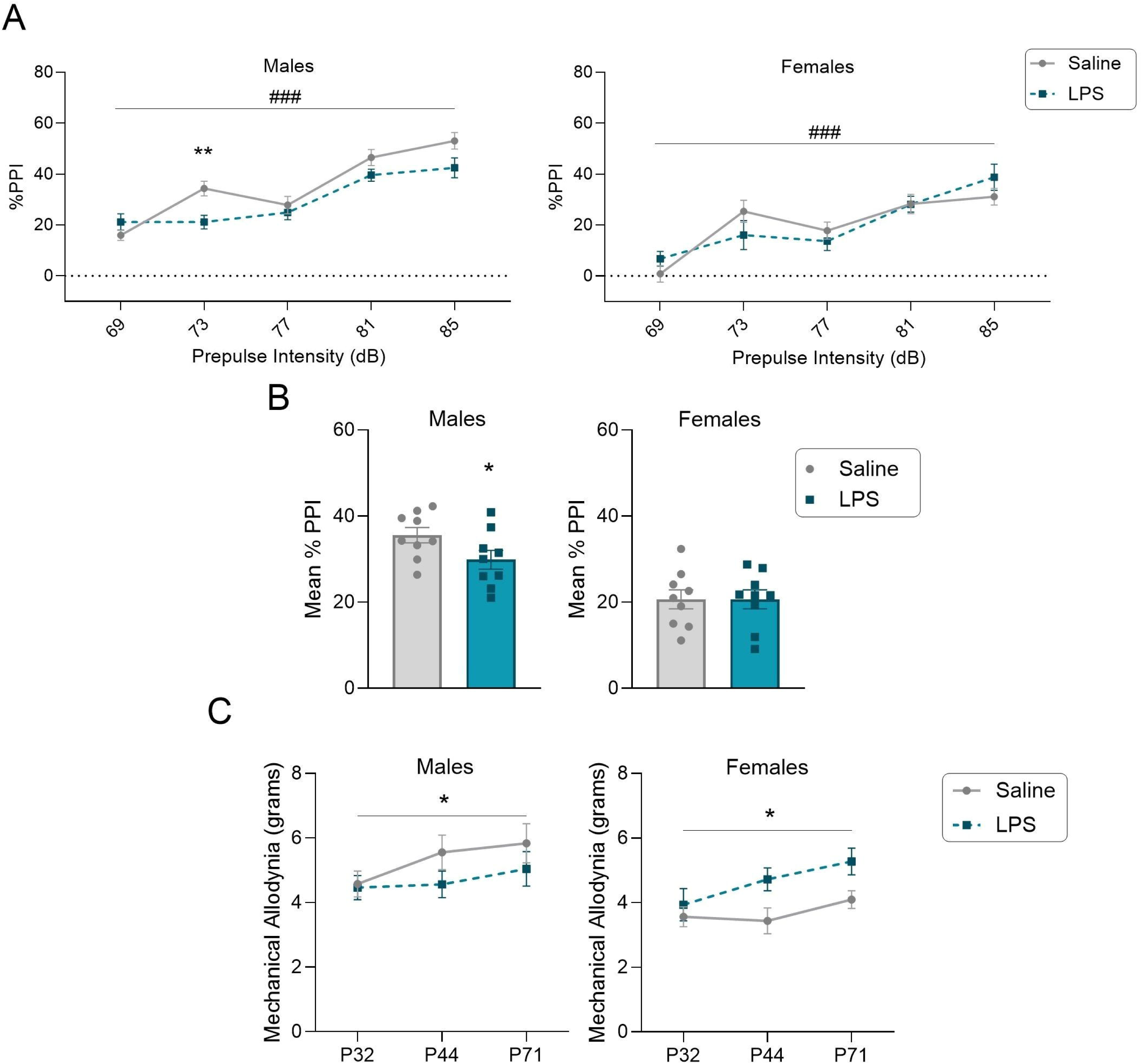
Adult offspring behaviors were affected by lactational maternal immune activation (MIA). Male and female offspring A) percent prepulse inhibition (%PPI) and B) mean %PPI of the acoustic startle response. C) Lactational MIA decreased mechanical allodynia thresholds (grams) in males (left-side) while increasing it in females (right-side) in the von Frey test. Saline: grey circles; lipopolysaccharide (LPS): blue squares. Data expressed as mean ± SEM, n = 9 litters/group. ###p<0.001, main effect of dB level; *p<0.05; **p<0.01; ***p<0.001, Saline versus LPS.

### Lactational MIA resulted in sustained alterations in immune function

Two-way ANOVA analysis revealed a significant interaction between P8 and adult LPS challenge on total leukocytes (males: F(1, 19) = 6.569, p = 0.019, *np^2^* = 0.849; females: F(1, 19) = 8.874, p = 0.040, *np^2^* = 0.203; **Figure 5A**). Male LPS-Saline animals had significantly higher leukocyte numbers than Saline-Saline (t(9) = 2.363, p = 0.042) and LPS-LPS male mice (t(9) = - 8.931, p = 0.001). However, despite the higher leukocyte counts at baseline, the P8 MIA male responses to a second adult LPS challenge did not differ from controls. Indeed, leukocyte numbers were reduced in both male (t(10) = −5.627, p = 0.001) and female (t(10) = −3.898, p = 0.003) Saline-LPS mice compared to Saline-Saline. Interestingly, female LPS-Saline mice had significantly reduced leukocyte numbers compared to their Saline-Saline female controls (t(9) = - 2.544, p = 0.031). Moreover, female LPS-Saline mice did not differ from LPS-LPS females (p>0.05), suggesting the lower baseline leukocyte counts in these P8 MIA offspring rendered them non-responsive to future LPS challenges, at least in terms of total leukocyte numbers.

**Figure 5.**
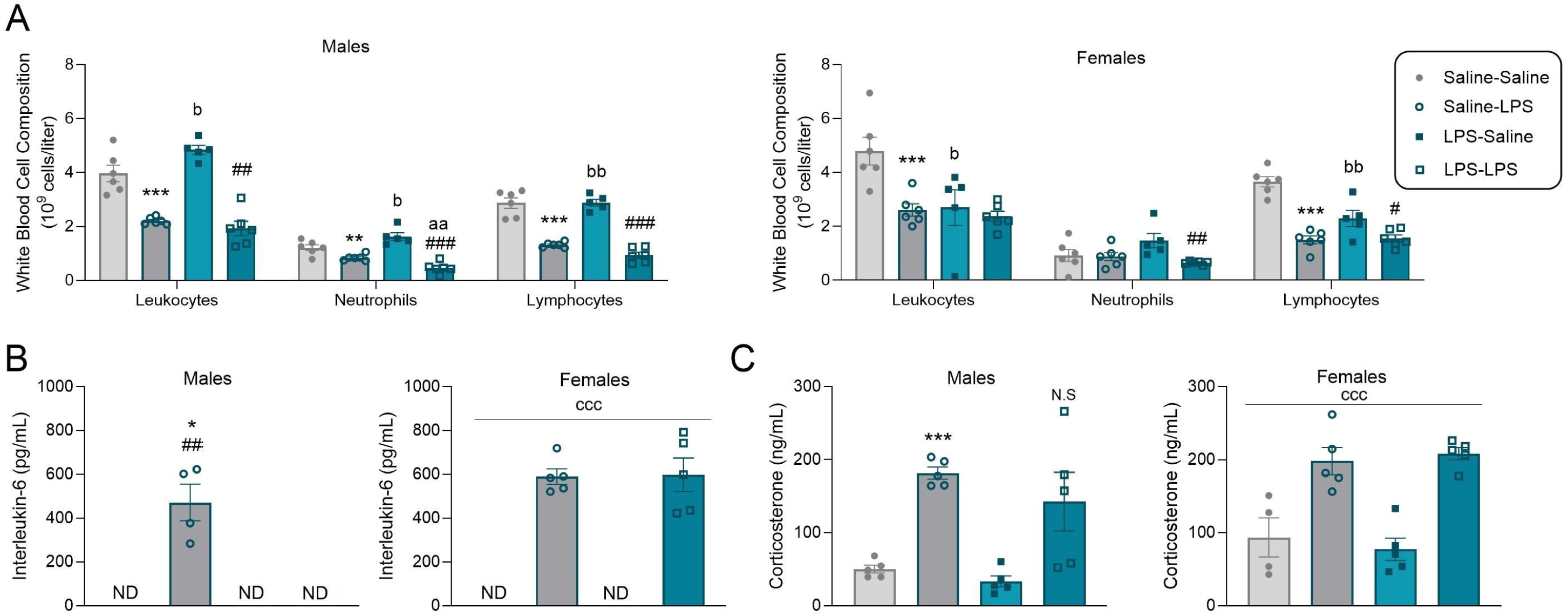
Adult offspring have altered immune functioning following lactational maternal immune activation (MIA). Plasma markers (n = 5-6 litters/group) for A) the number of white blood cells (leukocytes) measured, the absolute number of leukocytes that were neutrophils or lymphocytes (10^9^ cells/liter) directly measured, and B) interleukin (IL)-6 (pg/mL) demonstrating programming effects following the early life inflammatory challenge; one male Saline-LPS animal had undetectable levels of IL-6. C) Plasma corticosterone concentrations (ng/mL; n = 5 litters/group; one female Saline-Saline animal had undetectable levels of this hormone). Saline-Saline: solid circles and light grey shaded bars; Saline-LPS: open circles and solid dark grey bars; LPS-Saline: solid blue squares and light blue shaded bars; LPS-LPS: open squares and dark blue solid bars; lipopolysaccharide (LPS). Data expressed as mean ± SEM, Males on left side and females on right. *p<0.05, **p<0.01, ***p<0.001, Saline-Saline versus Saline-LPS; #p<0.05, ##p<0.01, ###p<0.001, LPS-Saline versus LPS-LPS; ^a^p<0.05, Saline-LPS versus LPS-LPS; ^b^p<0.05, Saline-Saline versus LPS-Saline; ^c^p<0.05, ^cc^p<0.01, ^ccc^p<0.001, main effect of adult LPS challenge.

There was a significant two-way ANOVA interaction for neutrophil concentrations in males (F(1,19) = 16.643, p = 0.001, *np^2^* = 0.467; **Figure 5A**). Similar to the leukocyte data, male LPS-Saline mice had significantly higher neutrophil concentrations than Saline-Saline males (t(9) = 2.83, p = 0.041) but here, P8 MIA male responses to a second LPS challenge differed from controls. Male Saline-LPS offspring had significantly lower levels of neutrophils compared to Saline-Saline (t(10) = −3.196, p = 0.010) and LPS-LPS males had reduced concentrations compared to LPS-Saline (t(9) = −7.433, p = 0.001) and Saline-LPS male mice (t(10) = −3.921, p = 0.003). In female animals, there was a main effect of adult LPS treatment on neutrophils (F(1,19) = 10.096, p = 0.005, *np^2^* = 0.347; **Figure 5A**). LPS-LPS females had significantly lower levels of neutrophils compared to LPS-Saline females (t(9) = −3.383, p = 0.008), but no other group comparisons differed (p>0.05). This suggests that P8 MIA exposure in females led to a slightly exaggerated neutrophil reduction in response to a second LPS challenge in adulthood.

A main effect of adult treatment was found for male concentrations of lymphocytes (F(1,19) = 178.320, p = 0.001, *np^2^* = 0.904; **Figure 5A**). Saline-LPS males had lower levels than Saline-Saline males (t(10) = −7.813, p = 0.001) and LPS-LPS males were lower than LPS-Saline (t(9) = −11.699, p = 0.001) and Saline-LPS male mice (t(10) = −3.245, p = 0.009). With respect to female animals, there was a significant interaction for lymphocytes between the lactational MIA and adult LPS treatment groups (F(1,19) = 13.453, p = 0.002, *np^2^* = 0.415; **Figure 5A**). Specifically, Saline-LPS (t(10) = −9.001, p = 0.001) and LPS-Saline (t(9) = −3.974, p = 0.003) animals had lower levels than Saline-Saline. This suggests sustained impacts of lactational MIA on the LPS-Saline females. Interestingly, LPS-LPS females had significantly lower lymphocyte concentrations compared to these LPS-Saline animals (t(9) = −3.383, p = 0.038), suggesting an exaggerated impact on lymphocyte levels following a second LPS hit.

A main effect of adult treatment was revealed for male (F(1,19) = 10.951, p = 0.004, *np^2^* = 0.366) and female (F(1,19) = 10.951, p = 0.004, n2 = 0.366) monocyte concentrations (**Supplemental Figure 1C,D**). While follow up tests did not confirm this effect in males, for females LPS-LPS animals had significantly reduced levels compared to LPS-Saline (t(9) = - 3.467, p = 0.007).

Eosinophils were associated with a main effect of lactational MIA in males F(1,19) = 5.790, p = 0.026, *np^2^* = 0.234; **Supplemental Figure 1C**) which was not confirmed by post hoc testing. Female data revealed a significant interaction between lactational MIA and adult LPS (F(1,19) = 12.199, p = 0.003, *np^2^* = 0.404; **Supplemental Figure 1D**). Saline-LPS females had higher eosinophil concentrations compared to LPS-Saline (t(9) = 4.229, p = 0.001) and LPS-LPS (t(9) = −3.268, p = 0.005) animals suggesting long term programming impacts following MIA that can be remodeled following a second LPS challenge in adulthood.

While basophils were not altered in males (p>0.05), in females there was a main effect of lactational MIA (F(1,19) = 4.382, p = 0.050, *np^2^* = 0.187; **Supplemental Figure 1C,D**). However, follow-up tests failed to identify specific group differences (p>0.05).

A two-way ANOVA revealed a lactational MIA by adult LPS interaction for plasma IL-6 in males (F(1, 16) = 10.865, p = 0.005, *np^2^*= 0.404; **Figure 5B**) and a main effect of adult LPS treatment in females (F(1,16) = 201.471, p = 0.001, *np^2^* = 0.926). For females, adult LPS challenge led to increased concentrations of plasma IL-6 (p = 0.001). On the other hand, in males, there was a clear developmental influence in that Saline-LPS animals had elevated IL-6 concentrations compared to Saline-Saline (t(7) = 2.907, p = 0.023), however, LPS-LPS animals failed to mount an IL-6 response to the adult inflammatory challenge (LPS-Saline versus LPS-LPS: p>0.05; Saline-LPS versus LPS-LPS: t(8) = −3.296, p = 0.011; one male Saline-LPS animal also failed to mount an IL-6 response).

With respect to plasma corticosterone, both males (F(1,16) = 32.895, p = 0.001, *np^2^* = 0.673) and females (F(1,15) = 45.292, p = 0.001, *np^2^* = 0.751;**Figure 5C**) had a significant main effect of adult LPS treatment, where corticosterone increased in response to the inflammatory stressor (males and females: p = 0.001). Given the attenuation of the IL-6 response in male LPS-LPS animals, we conducted secondary analyses to more clearly determine if the corticosterone response was also altered in LPS-LPS animals, similar to other neonatal immune models (Shanks et al., 2000; Shanks et al., 1995; Walker et al., 2009; Hodgson et al., 2001; Ellis et al., 2005).As expected, male Saline-LPS mice had higher levels of this stress hormone compared to Saline-Saline animals (t(8) = 13.369, p = 0.001). However, secondary analyses showed that LPS-LPS animals were not different than LPS-Saline or Saline-LPS animals, suggesting modest attenuation of this hormone.

## Discussion

Early-life immune challenges can leave lasting imprints on offspring brain and behavior, with some consequences emerging only after a second stressor later in life. The present study provides a cross-species validation of the lactational maternal immune activation (MIA) LPS model, supporting its use for investigating the mechanisms of these effects. Offspring outcomes following lactational MIA have been primarily evaluated in rats (Vilela et al., 2013; Nascimento et al., 2015; DeRosa et al., 2022) and adapted here for C57BL/6J mice; the replication and extension of key offspring behavioral and immunological phenotypes underscores the translational relevance and reproducibility of this early life model of maternal infection. Importantly, our findings align with a growing body of literature suggesting that inflammatory insults during critical windows of early development can elicit lasting consequences on offspring physiology and behavior, extending into adulthood (Knuesel et al., 2014; Bilbo & Schwarz, 2012; Kentner et al., 2019; Gumusoglu et al., 2019; Cattane et al., 2022; Otero & Antonson, 2022).

Central to this study was the observation that a single inflammatory event during the lactational period, when restricted to the dam, was sufficient to alter developmental trajectories in offspring, both behaviorally and in their capacity to respond to a subsequent immune challenge. With respect to behavior, while lactational MIA in mouse dams elicited expected and robust sickness behaviors within hours of the immune challenge, its impact on maternal care (indexed via nest construction quality) was modest and transient. This subtle disruption is in contrast to findings in some prenatal (Ronovsky et al., 2017; Zambon et al., 2024) and lactational (Vilela et al., 2013) MIA studies where sustained impairments in maternal behavior have been reported yet align with other lactational MIA data where maternal care was not adversely altered (Aubert et al., 1997; DeRosa et al., 2022; Nascimento et al., 2015). These findings suggest that while both prenatal and postnatal immune activation elicit maternal sickness behaviors, the timing, type- and/or dose of the immunogen may differentially affect caregiving, possibly due to differences in hormonal profiles at the point of challenge.

Despite the relatively preserved maternal care, our results highlight that certain consequences of MIA are not immediately apparent but may manifest later in development. While neonatal behavior was not significantly impacted by lactational MIA, consistent with reports in some prenatal MIA models where early reflexive behaviors remain intact (e.g., Malkova et al., 2012; Staley et al., 2017), downstream effects were evident in both juvenile and adult offspring. One limitation of our study is that multiple behavioral tests were run on P10, which may have stressed the neonates, masking group differences. This may account for the inconsistencies in huddling behaviors displayed between the rat and mouse lactational MIA models in particular. At the juvenile age, lactational MIA accelerated puberty in female offspring, a finding also observed in prenatal MIA (Zhao et al., 2022) and other animal models of early-life stress (Grassi-Oliveira et al., 2016; Granata et al., 2024). However, this acceleration was quite modest, only advancing puberty by about one day in the present study.

In adulthood, male (but not female) MIA offspring exhibited reduced prepulse inhibition (PPI) of the acoustic startle response, a commonly used measure of sensorimotor gating. This replicates findings from both prenatal and postnatal MIA models in rats and mice (Herrero et al., 2023; Richetto et al., 2017; Zhao et al., 2022; DeRosa et al., 2022; Kentner et al., 2019). Moreover, the enhanced social preference in juvenile males and reduced social preference in females, both of which were recovered by adulthood, support findings that 1) immune challenges likely lead to divergent outcomes through interactions with factors such as the environment, offspring sex, species, and timing of infection (Schaer et al., 2024; Liu et al., 2024; Connors et al., 2014; Kentner et al., 2019; Meyer et al., 2009), and 2) the trajectory of certain vulnerabilities that emerge following MIA can unfold and/or recover across developmental windows (MacRae et al., 2015b; Dinel et al., 2014; Zuckerman et al., 2013; Meyer et al., 2006; Paylor et al., 2016). A summary table outlining the known effects of lactational MIA in rats and mice can be found in **Table 1**, identifying some remaining areas for future research. Collectively, these findings emphasize that the lactational period is a sensitive window where maternal communication signals can shape offspring behavior, given that postnatal MIA recapitulated several key features of the prenatal MIA models (Estes & McAllister, 2016; Kentner et al., 2019; Meyers et al., 2009). This was in addition to also revealing unique effects related to the timing and context of inflammation **Table 1**).

**Table 1.**
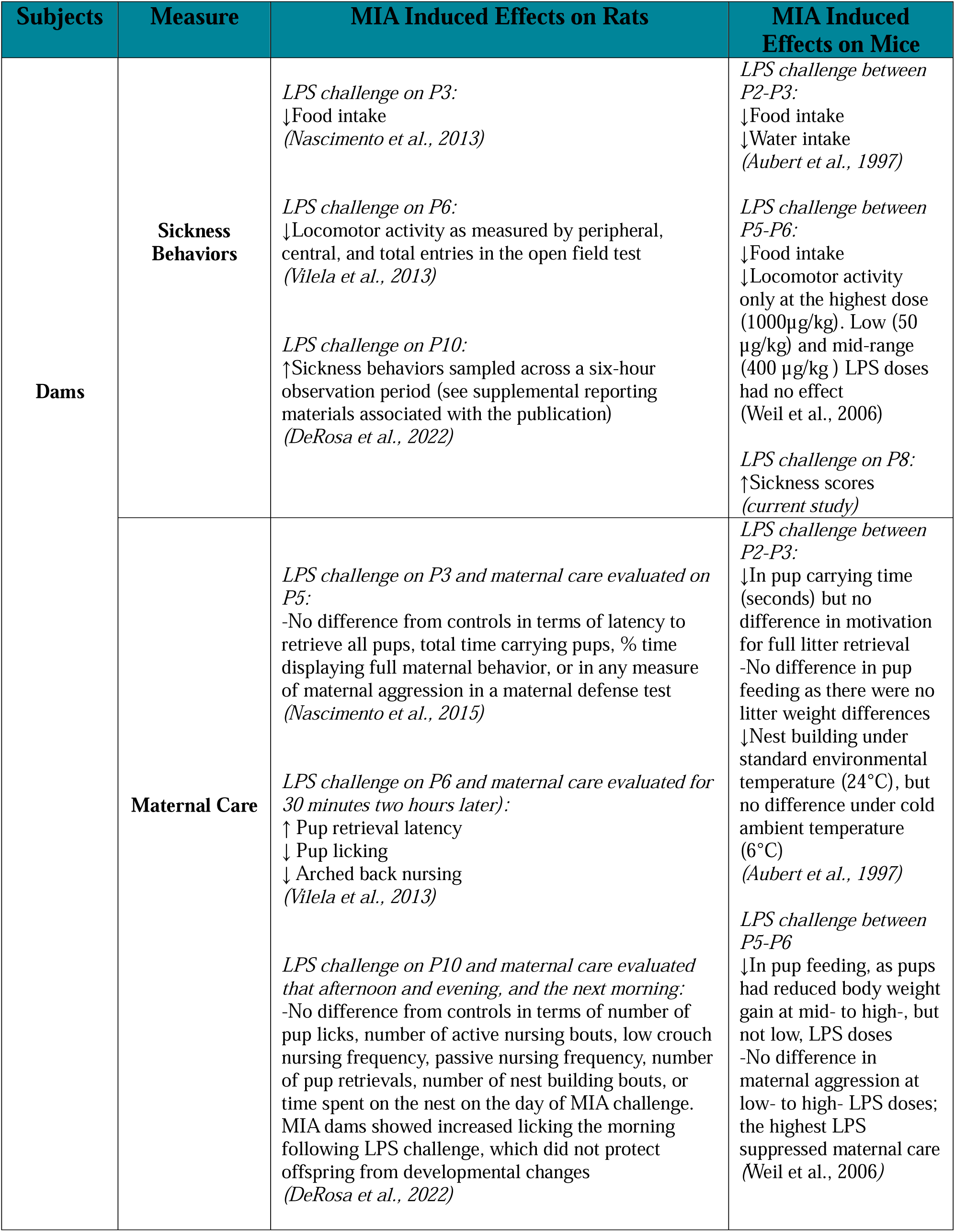
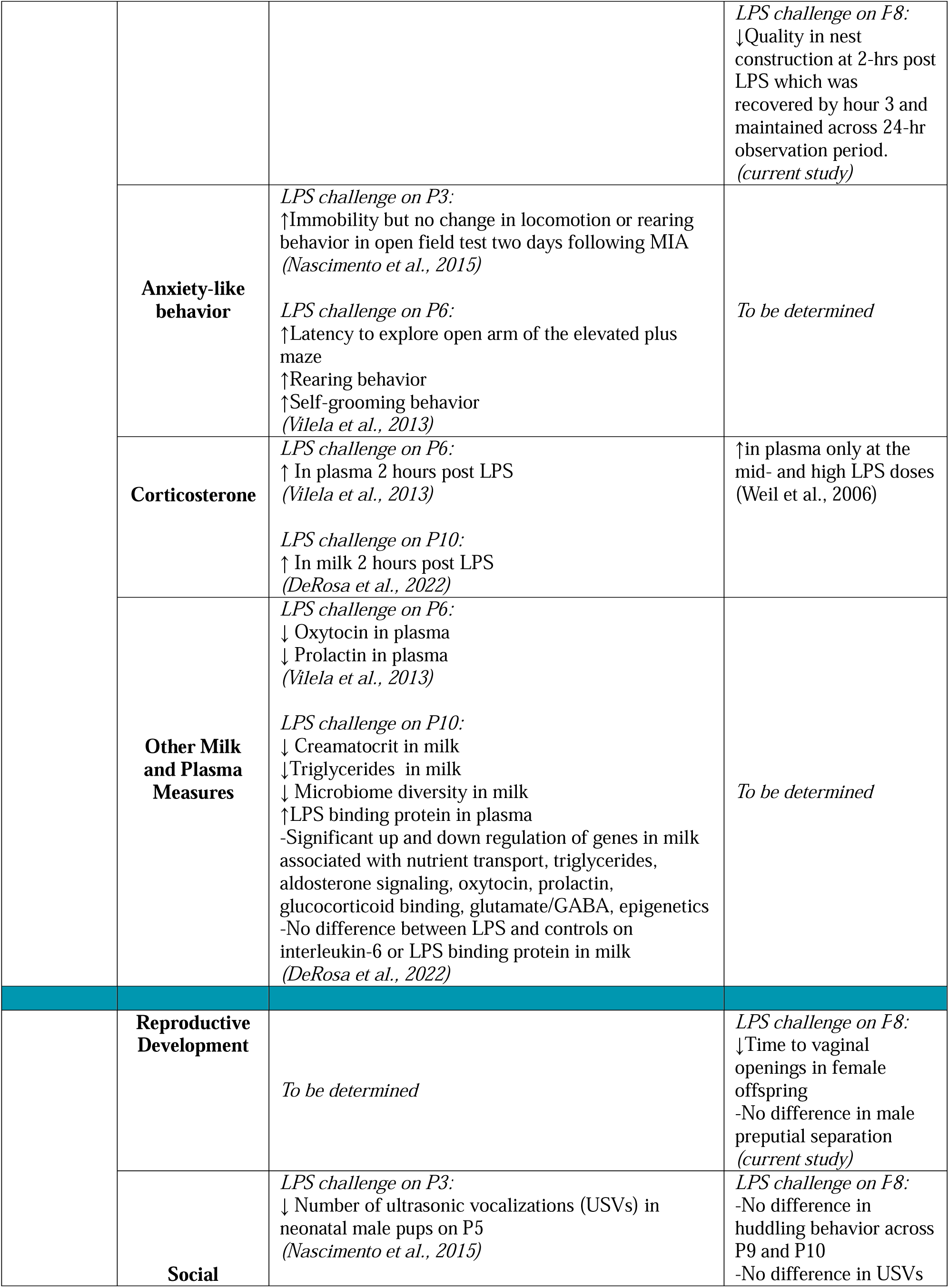
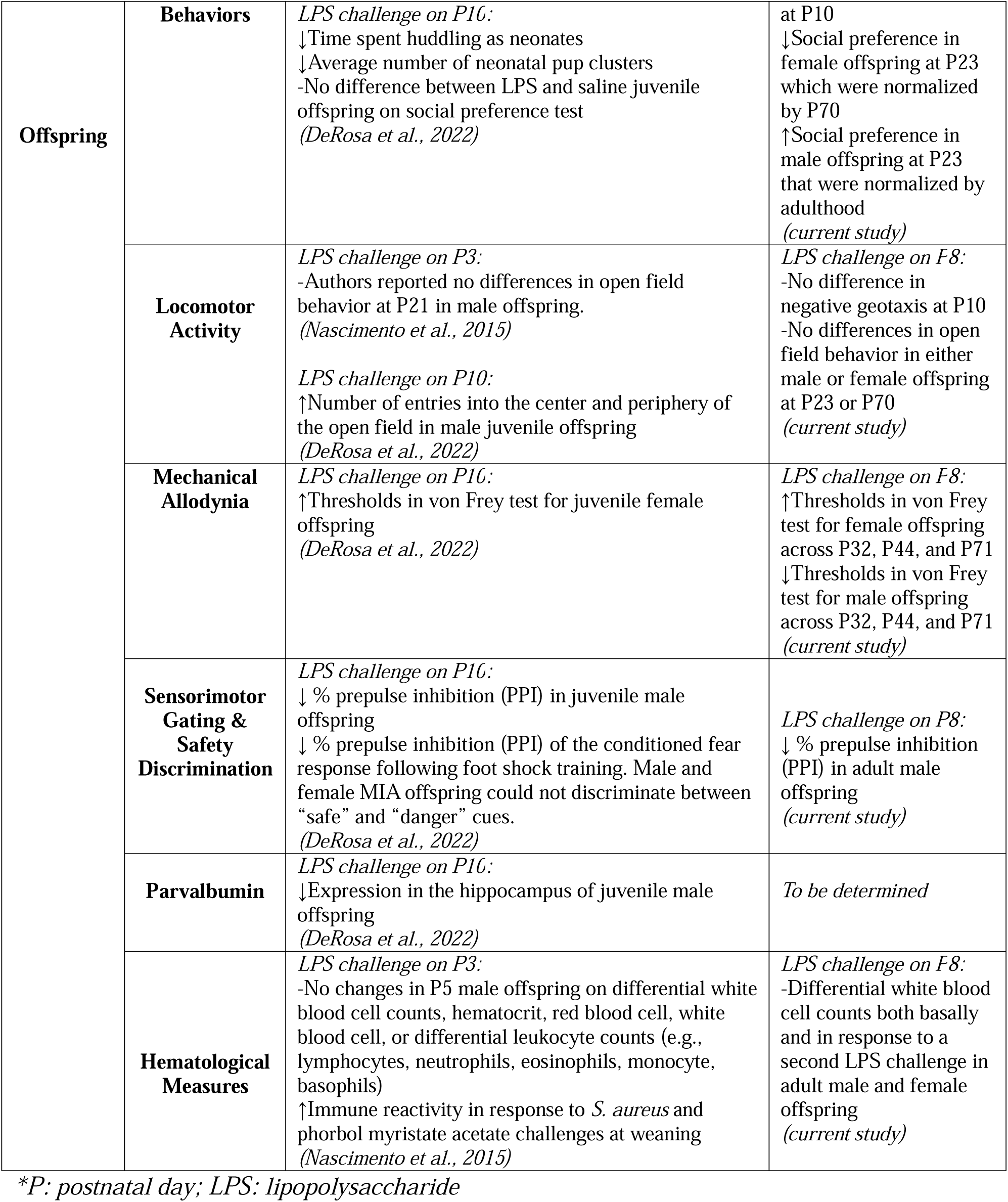
Summary table comparing some known and unknown effects of maternal immune activation (MIA) versus saline on mice and rats.

White blood cells (WBC) are essential for immune surveillance and inflammatory responses (Janeway et al., 2001). However, their functionality and reactivity are not fixed; throughout life they are shaped by developmental cues and environmental exposures (Palomino-Segura et al., 2023; Sailhamer et al., 2010; Nascimento et al., 2015; Netea et al., 2016). In our animals, an acute adult LPS challenge resulted in general reductions of neutrophil and lymphocyte numbers (Saline-LPS versus Saline-Saline animals), which can be indicative of a bacterial or viral infection (El Brihi & Pathak, 2024). Our data also demonstrates that lactational MIA may exert enduring immunomodulatory effects, as evidenced by altered levels of basal WBC populations in adulthood. While previous work in rats showed no changes to WBC counts in P5 neonatal offspring (following a P3 lactational MIA challenge), MIA males demonstrated higher immune reactivity at weaning (Nascimento et al., 2015). Here, there was an elevation in lymphocyte and neutrophil levels in adult LPS-Saline mice (compared to Saline-Saline), suggesting that lactational MIA exposure lead to dysregulation in WBC concentrations. Notably, elevations in WBCs are associated with atherosclerosis (Kabat et al., 2017), which could indicate an increased risk for cardiovascular disease in these animals. Moreover, neutrophil and cytokine responses (e.g., IL-6), after the second LPS challenge, reflected long-term programming of both the innate and adaptive arms of the immune system in male offspring. Specifically, both neutrophils and IL-6 levels were attenuated in LPS-LPS versus Saline-LPS mice. Female neonatal LPS mice had reduced concentrations of leukocytes at baseline (LPS-Saline versus Saline-Saline), that did not respond further following a second LPS challenge (LPS-Saline versus LPS-LPS). Low leukocyte concentrations may be reflective of a reduction in their production, higher rates of utilization, or even destruction of these cells (El Brihi & Pathak, 2024). However, these distinct immune profiles in offspring exposed to lactational MIA do not appear to be maladaptive as the magnitude of other immune response measures did not generally differ between neonatal LPS and Saline exposed controls. That said, immune responses could vary depending on the type of immune exposure; for example, whether the second hit is a bacterial, viral, or fungal challenge, in addition to what the level or magnitude of that exposure is.

Male offspring in particular demonstrated aberrant responses across a greater number of the immune parameters evaluated, supporting the concept of sex-specific programming of immune function (Klein & Flanagan, 2016), which future studies can follow up on more specifically. As noted, exposure to LPS in adulthood elicited robust plasma IL-6, in addition to corticosterone, responses when raised by mothers treated with Saline during the lactational period (e.g., Saline-LPS versus Saline-Saline males). These effects were attenuated in males raised by MIA dams (e.g., LPS-LPS versus LPS-Saline). Observations of reduced circulating cytokine responses to a second LPS challenge have been reported in adult male rats that had directly received LPS in the neonatal period (Ellis et al., 2005; Kentner et al., 2010). With respect to plasma corticosterone, the LPS-induced attenuation was quite modest in our lactational MIA model (as evaluated via secondary follow up tests), and the variability in the data suggests subpopulations of resilient versus susceptible animals in the LPS-LPS group. Attenuation of plasma corticosterone release has previously been observed when the “second hit” in adulthood was a different, or “heterotypic” stressor-type from the neonatal LPS challenge (Shanks et al., 2000; Shanks et al., 1995; Walker et al., 2009). In general, the context of the subsequent challenge is likely key as some studies report exaggerated corticosterone responses to stressors (Hodgson et al., 2001; Ellis et al., 2005), especially in response to a “homotypic” second challenge with LPS in male animals (Ellis et al., 2005). This latter effect is thought to be mediated via TLR4-and COX-2 associated mechanisms in the two-hit neonatal LPS model (Spencer et al., 2011; Mouihate et al., 2010).

Most pathogens are not passed through breastmilk to nursing infants, so rates of mother to child transmission of infections are low (Yeo et al., 2022;CDC, 2025). Rather, breastfeeding is recognized to provide protection to babies from illness, through antibodies and other milk borne factors, and is therefore recommended during most maternal conditions (Yeo et al., 2022; World Health Organization, 2009, CDC, 2025). Depending on the infection, masks and other safety precautions may also be recommended (Wijenayake et al., 2023; Yeo et al., 2022). In our previous lactational MIA work with rats (DeRosa et al., 2022), we showed that LPS binding protein, a direct measure of LPS (Citronberg et al., 2016; Bishara et al., 2012), also did not cross from milk to nursing pups. Specifically, LPS binding protein was significantly higher in the plasma of MIA exposed rat dams compared to saline, but not in the milk where LPS binding protein was virtually undetectable (DeRosa et al., 2022). Therefore, unlike the neonatal two-hit rat models (Ellis et al., 2005; Kentner et al., 2010; Spencer et al., 2011; Shanks et al., 2000; Shanks et al., 1995; Walker et al., 2009), it is unlikely that lactational MIA offspring were directly exposed to the immunogen. This underscores the potential for another indirect maternal communication cue (e.g., a constituent in milk; Wijenayake et al., 2023) to act as a “heterotypic” first stressor, shaping future immune responses to an adult LPS exposure. Given the relatively short life span of circulating leukocytes such as lymphocytes, and especially neutrophils and monocytes (Palomino-Segura et al., 2023; Netea et al., 2016), the programming effects of WBCs are likely taking place via modulation of hematopoietic stem and progenitor cells and epigenetic rewiring (Mitroulis et al., 2018; Netea et al., 2016). While the specific mechanism(s) by which early life inflammatory stressors cause changes to adult immune functioning appears to depend on the animal model investigated, the overall consequence of altered immunomodulation appears consistent across these developmental challenges.

Overall, the translational value of this work lies in replicating, and extending upon, the behavioral and immune effects of lactational MIA from rats to mice (see **Table 1**). While mice and rats share broad physiological and developmental similarities, they also have notable species-specific differences in both immune and behavioral repertoires. Therefore, demonstrating the cross-species applicability of the lactational MIA model strengthens its utility in preclinical research. Our findings also emphasize the need for additional studies that assess not only behavioral phenotypes across species, as was the focus here, but also on peripheral immune signatures, which may serve as early biomarkers of risk or resilience following early life adversity. This work has implications for understanding how maternal inflammation during nursing, not just pregnancy, can shape offspring behavioral and immunological trajectories.

## Acknowledgements

This project was funded by NIMH under Award Number R15MH114035 (to ACK), the Massachusetts College of Pharmacy and Health Sciences (MCPHS) Center for Research and Discovery (JM), and a Massachusetts Life Sciences Center Capital Grant. Figure 1 was made with BioRender.com. The content is solely the responsibility of the authors and does not necessarily represent the official views of any of the financial supporters.

## CRediT authorship contribution statement

**Jailyn A Merengueli**: Investigation, Data curation, Formal analysis, Writing – review & editing, **Amanda C. Kentner:** Conceptualization, Study Design, Investigation, Data curation, Formal Analysis, Writing – original draft, review & editing, Resources, Funding acquisition.

---

**Supplemental Figure 1.** Adult male and female offspring A) social preference index and B) percent time spent in center of the open field. Saline: grey circles; lipopolysaccharide (LPS): blue squares (n = 9 litters/group). Second hit challenge data for the absolute number of white blood cells (leukocytes) that were monocytes, eosinophils, or basophils directly measured (10^9^ cells/liter) for C) male and D) female offspring.). Saline-Saline: solid circles and light grey shaded bars; Saline-LPS: open circles and solid dark grey bars; LPS-Saline: solid blue squares and light blue shaded bars; LPS-LPS: open squares and dark blue solid bars; Data expressed as mean ± SEM. *p<0.05, **p<0.01, ***p<0.001, Saline-Saline versus Saline-LPS; #p<0.05, ##p<0.01, ###p<0.001, LPS-Saline versus LPS-LPS; ^a^p<0.05, Saline-LPS versus LPS-LPS; ^b^p<0.05, Saline-Saline versus LPS-Saline.

