## Supplementary figures and images for "Maternal immune activation during the lactational period alters offspring behavior, reproductive development, and immune function in mice"

### Supplemental Figure 1

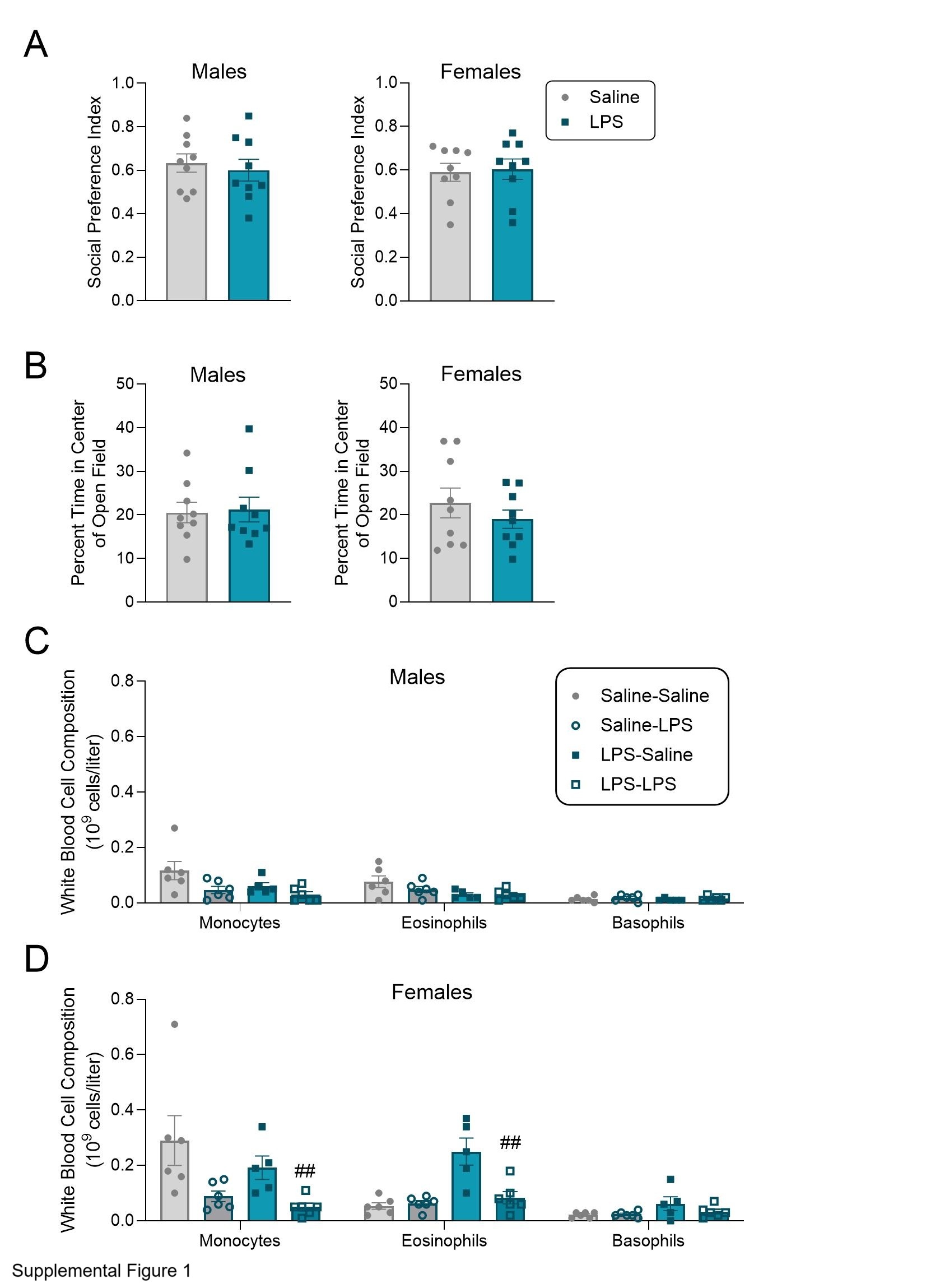
